# Fast and robust objective EEG audiometry

**DOI:** 10.1101/2024.05.22.595285

**Authors:** T. Guyonnet-Hencke, O. Portoles, M. de Vries, E. Koderman, A. Winkler, J. Goodall, P. Desain, J. Thielen, M. Schulte, A. J. Beynon, M. van Kesteren

**Affiliations:** MindAffect B.V., Netherlands; Donders Institute for Brain, Cognition and Behaviour, Radboud University, Netherlands; Hörzentrum Oldenburg gGmbH, Germany; Center for Brain and Senses, Radboud University Medical Center, Netherlands

**Author notes:** shared co-authorship. shared co-authorship and contact author.

## Abstract

The current ‘gold standard’ of audiometry relies on subjective behavioral responses, which is impractical and unreliable for certain groups such as children, individuals with severe disabilities, or the disabled elderly. This study presents a novel electroencephalography (EEG) system that is easy to setup and estimates audiometric thresholds quickly. Air-conduction audiometric thresholds at 250, 500, 1000, 2000, 4000, and 8000 Hz and 5 dB resolution were estimated from ten elderly patients with asymmetric sensorineural hearing loss and five normal hearing young adults using three different systems: the novel EEG system, conventional pure-tone audiometry (PTA), and an automated behavioral test with the same stimulus properties as in the EEG test. EEG data was collected for 15 minutes from 32 semi-dry EEG electrodes. Later, the EEG system was trimmed to 8 electrodes and 7.5 minutes of data with satisfactory results. Correlation and regression analysis validated the hearing thresholds derived from both EEG configurations relative to the behavioral hearing thresholds–Pearson’s correlation of 0.82 between PTA and 8-electrode 7.5-minute EEG data. The results of this study open the door to fast and objective hearing threshold estimation with EEG.

## 1 Introduction

Hearing loss is a significant public health concern affecting a considerable portion of the global population, with approximately 20% of individuals worldwide experiencing some degree of hearing impairment. Among children, 60% of hearing loss cases are attributed to avoidable causes that could be addressed by implementing preventive measures (World Health Organization, n.d.). The consequences of untreated hearing loss extend beyond auditory impairment, impacting various aspects of individuals’ lives, including communication, social interaction, and overall quality of life (Nordvik et al., 2018). Furthermore, recent studies have highlighted the association between untreated hearing loss in midlife and an increased risk of dementia, making it a critical modifiable risk factor for cognitive decline (Livingston et al., 2020).

The detection and diagnosis of hearing loss play a crucial role in preventing its adverse effects and facilitating timely interventions. The gold standard for diagnosing hearing loss is pure-tone audiometry (PTA). PTA is a behavioral measurement of audiometric thresholds that relies on a yes/no response of the participant to the perception of a pure-tone. However, the reliability of PTA may be compromised by various factors including difficulties in understanding the instructions, lack of cooperation, or potential malingering (Wadhera et al., 2017; Swami & Kumar, 2019). This impracticality is particularly common when testing hard-to-test individuals such as infants, children, people with disabilities, and the elderly (Ferguson et al., 2023; Tarawneh et al., 2022). It is reported that approximately 40% of adults with dementia have difficulties completing PTA (Anthea et al., 2019). Young children can have difficulties engaging in behavioral tasks. For some children, the gap between their developmental and chronological age makes it difficult for them to complete age-appropriate behavioral hearing tests (Teplitzky et al., 2019). Additionally, Widen (1993) suggested that the supplementary information added by objective testing in children, such as the type of hearing loss, must not be disregarded. These limitations underscore the need for easily accessible, inclusive, and reliable auditory assessment tools that can accommodate diverse patient populations, and be administered by healthcare professionals without specialized audiological training (Ferguson et al., 2023).

Objective audiometric methods that do not necessitate active patient engagement are available as alternatives to conventional behavioral audiometry. One alternative technique to standard behavioral audiometry is otoacoustic emission (OAE). OAE measures outer hair cells’ integrity through sound delivery. However, measurement of otoacoustic emission cannot provide accurate hearing thresholds and degree of deafness as the movement of hair cells is not informative enough for a precise hearing threshold estimation. Moreover, OAEs traveling through the middle ear can be affected by middle ear diseases. Thus, OAE is not sufficient to differentiate between conductive and sensorineural hearing loss (Kemp et al., 1986; Probst et al., 1991). Another alternative to conventional behavioral audiometry is electrophysiological methods such as auditory brainstem response (ABR) or the auditory steady-state response (ASSR). ABR and ASSR can objectively predict audiometric thresholds for a limited frequency range, usually 500 to 4000 Hz. However, ABR and ASSR require 20–25 or 32–60 min to estimate 8 thresholds respectively (Sininger et al., 2018), and they are very vulnerable to artifacts. The measurement of electrophysiological responses to auditory stimuli is highly susceptible to noise and artifacts from exogenous and endogenous sources. Consequently, it is important to maintain the patient calm and still for large periods of time to maximize the signal-to-noise ratio and mitigate the likelihood of inaccuracies in the threshold estimation.

Children represent another population for which behavioral testing can be challenging and methods such as ABR and ASSR are therefore needed. However, these tests often require the child to be asleep (Sininger et al., 2018) to minimize the influence of artifacts. Sedation is often required for patients who cannot cooperate in this regard, such as young children (Sininger et al., 2018; Janssen et al., 2010). Thus, the amount of thresholds that can be tested is dependent on the duration of the sleep or patient collaboration.

We propose a novel objective audiometric technique that uses short electroencephalographic (EEG) recordings (7.5 min or less) with fast and short auditory evoked potentials (AEP) that are coded to mitigate the challenges in current audiometric tests. First, EEG audiometry removes the subjectiveness and inaccuracy of behavioral responses by not requiring active patient collaboration nor the interpretation of behavioral signs by technicians. Second, the proposed EEG system mitigates the sensitivity to noise and artifacts characteristic of current EEG methods which require long recording sessions to enhance the signal-to-noise ratio. The technology powering the proposed EEG method suppresses the influence of artifacts to improve the signal-to-noise ratio of AEPs and reduce the recording times which makes it less vulnerable to patient’s fatigue, lack of cooperation, and artifacts compared with ASSR or ABR. Third, a key component of the proposed EEG system is the use of short stimuli with a fast presentation rate which also contributes to further reduction in the duration of an EEG audiometric session.

The first goal of this study was to assess the validity of the hearing thresholds estimated from 15 minutes of 32-channel EEG data with respect to PTA. To this end, the EEG data was analyzed blindly without information about behavioral hearing thresholds of the participants. Next, the feasibility of a trimmed EEG system with fewer channels and less EEG data was assessed. Subsequently, a trimmed EEG system with 8 channels and 7.5 minutes of EEG data was validated with PTA and an automated behavioral test with the same stimulus properties as in the EEG test (autoBEH). Finally, the systematic offsets between EEG, PTA, and autoBEH hearing thresholds were analyzed.

## 2 Methods

### 2.1 Participants

For this study, 15 participants were tested by Hörzentrum Oldenburg (6 females and 9 males), of whom 10 were hearing impaired (group HI, mean age of 70 ± 7 years) and 5 were control young-adult participants with no hearing impairment (group CTRL, mean age of 30 ± 5 years; 4 of them were employees of MindAffect). HI participants exhibited sensorineural hearing loss and an asymmetry in hearing: one ear had a more pronounced impairment than the other one. For 8 out of the 10 HI participants, the more damaged ear corresponded to the right ear. Ethical approval was granted by the ethics committee (Kommission für Forschungsfolgenabschätzung und Ethik) of the Carl von Ossietzky University in Oldenburg, Germany (Drs.EK/2021/031-05). Tables S1 and S2 in the Supplementary Section shows the average PTA thresholds for each frequency for both groups.

### 2.2 PTA testing

Thresholds were estimated behaviorally using PTA with the Madsen Astera system and Sennheiser HDA200 circumaural headphones. Thresholds were estimated for 250, 500, 750, 1000, 1500, 2000, 3000, 4000, 6000, and 8000 Hz at both ears. For the following analyses, only thresholds at 250, 500, 1000, 2000, 4000, and 8000 Hz were considered, as only those frequencies were collected with the other tests of this study. The stimuli were pure-tone with a duration varying between 1 to 3 s, according to the British Society of Audiology (British Society of Audiology, n.d.). Stimuli were presented sequentially at various volumes, and participants were instructed to press a button to indicate that they had perceived the presented stimulus. Hearing thresholds were identified following the guidelines of the British Society of Audiology: after a correctly perceived stimulus, volume was decreased by 10 dB, and increased by 5 dB after a failure to perceive the stimulus (British Society of Audiology, n.d.). The volume of the stimuli was calibrated following the §14 Medizinprodukte Betreiberverordnung (MPBetreibV) protocol (Bundestag, 2002). The volume of PTA and EEG stimuli were calibrated the same day following the same protocol to ensure the same decibels hearing level (dB HL) between the two systems.

### 2.3 EEG testing

The proposed EEG system was a 32-channel EEG amplifier (Semi-dry Versatile, BitBrain, Spain) with semi-dry electrodes (i.e. humid sponges). The sampling rate was 256 Hz with 24-bit resolution. The electrodes were placed with a cap following the standard 10-20 system. The reference electrode was placed on FCz and the ground electrode on AFz. The EEG data was preprocessed by downsampling to 64 Hz with an anti-aliasing filter. A notch filter was applied at 50 Hz to remove any power-line interference and the data was subsequently bandpass filtered between 3 and 15 Hz using a sixth-order Butterworth filter. EEG data was epoched to 15 s, synchronized to the presented stimuli, resulting in a total of 60 epochs for 15 min of EEG recording. To remove any eye movement artifact, we used the signal-space projection technique to extract a projection space orthogonal to noise signals corresponding to eye movement artifacts. This was done by identifying windows with excessive power (2 standard deviations above the mean voltage power) within the 0.5-8 Hz frequency bandwidth in Fp1 and Fp2 and computing this projection space through singular value decomposition. The EEG data from every channel was then projected onto this subspace to remove eye-movement activity from the EEG signal (Uusitalo & Ilmoniemi, 1997).

Auditory stimuli were pure-tones with a duration of 30 ms for each of the tested frequencies: 250, 500, 1000, 2000, 4000, and 8000 Hz. Rise, fall, and plateau time of the pure-tones were all equal to 10 ms. The pure-tones were presented to the participants using Sennheiser DD-45 supra-aural headphones. Stimulus presentation was controlled by an HP Pavillon x360 laptop, and sounds were sent from the laptop to headphones through a FocusRite Scarlett 2i2 3rd-generation audio amplifier. The volume intensities ranged from 0 to 80 dB HL for the HI group and from -10 to 70 dB HL for the NH group, with steps of 5 dB, plus one silent stimulus (control level) resulting in a total of 18 levels. The volume range was shifted upwards for the HI group to better capture their audiometric thresholds while keeping the number of volumes tested constant. Stimuli were presented pseudo-randomly at an ear, volume, and frequency combination. Stimuli with a random combination of ear, frequency, and volume intensity were presented at a rate of 5 Hz. In total, 12 audiometric thresholds were measured for each participant, 6 by ear.

EEG data was recorded for a total duration of 15 min which on average presents approximately 20 times each of the 216 stimuli (i.e., 2 ears × 18 volumes × 6 frequencies). The EEG tests did not require active participation from the participants. The participants were asked to remain still to the extent possible to prevent any artifacts in the EEG data. To prevent tiredness and keep the participants calm, they watched a silent wildlife documentary video while auditory stimuli were presented. There was a fixation cross at the center of the screen to encourage the participants to fixate and avoid eye movements, to prevent contamination of the EEG.

### 2.4 autoBEH testing

A third testing condition was a behavioral audiometric testing similar to PTA but with the same stimulus properties as the EEG test (autoBEH). The autoBEH acted as a “control” condition, as many hardware components were not shared between the EEG and PTA systems. Components that differed between EEG and PTA were the headphones — circum-aural for the PTA testing and supra-aural for the EEG testing — and the duration of the stimuli — between 1 and 3 s for the PTA testing and 30 ms for the EEG testing. Circum-aural headphones cannot be used for the EEG system due to the overlap with the EEG electrodes, which can impair the signal recording at those electrodes. Short-duration stimuli were used for the EEG system to allow a fast presentation rate, a key feature of this system, contributing to its speed. Therefore, the autoBEH contributed to addressing whether the differences (if any) between the output of the PTA and EEG testing were due to the nature of the testing, behavioral versus electrophysiological, or the material used. As for the EEG testing, the autoBEH used an HP Pavillon x360 laptop to control the test which sent pure-tones of 30 ms to supra-aural Sennheiser DD45 through a FocusRite Scarlett 2i2 3rd-generation audio amplifier. This behavioral test was automatized with a modified Hughson-Westlake staircase procedure (Hughson & Westlake, 1944). The staircase procedure started from the higher volume intensity (70/80 dB HL) to the lowest (−10/0 dB HL). As in the Hughson-Westlake procedure, if a stimulus was successfully perceived the volume intensity of the next stimulus was decreased by 10 dB. If the stimulus was not perceived, then the volume intensity of the next stimulus was increased by 5 dB. Audiometric thresholds were estimated at 250, 500, 1000, 2000, 4000, and 8000 Hz in ascending order, for first the left ear and then the right ear. Participants answered by pressing the space bar of the laptop if they perceived the presented stimulus.

### 2.5 Experimental protocol

Each participant underwent auditory assessments through three distinct methods: PTA, autoBEH, and EEG testing. PTA testing was conducted first in one room followed by autoBEH and EEG testing in a separate room. The three conditions were done on the same day and both rooms were soundproof.

### 2.6 EEG analysis

The multi-channel EEG signals **X** of each participant were modeled as the combination of the AEP **r**, the AEP’s amplitude weights ***θ***, and the contribution to the AEP from each electrode **a**, as follows:

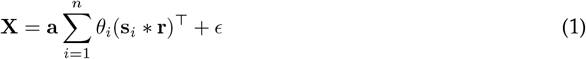

where the data **X** ∈ ℝ^*c×m*^ represents the EEG measurements across *c* channels for *m* samples, **a** ∈ ℝ^*c*^ denotes the spatial pattern associated with the modeled source across *c* channels, which illustrates how the neural activity from the source spreads across the scalp through the EEG channels. **s**_*i*_ is a vector of *m* samples representing rows, *i*, of the matrix **S** ∈ ℕ^*n×m*^ of *n* stimulus modalities over *m* samples, conveyed as a 0-1 indicator matrix or unit impulses, where *i* represents the modality of the stimulus and ranges from 1 to *n*. The number of stimulus modalities is *n* = *f × v*, where *f* is the number of auditory stimuli (here 12, 6 frequencies times 2 ears), and *v* is the number of volume intensities (here 18). The vector **r** ∈ ℝ^*τ*^ signifies the AEP, i.e., the electrical activity of the response to the isolated auditory stimulus persisting for *τ* samples. The asterisk *\** symbolizes the linear convolution of the time window of the event responses (*τ* samples) throughout the entire duration of the recording (*m* samples).

For this context of AEP analysis, we used *τ* corresponding to 400 ms, which at a sampling rate of 64 Hz corresponds to approximately 25 samples. The scalar *θ*_*i*_ ∈ ℝ represents the amplitude weighting of the AEP **r**, which can vary depending on the stimulus modality *i*. The term *ϵ* ∈ ℝ^*c×m*^ accounts for unmodeled signal and noise in the data.

The parameters **r** and **a** can be determined through a ‘reconvolution’ canonical correlation analysis (CCA) (Thielen et al., 2015, 2021). CCA is a technique exploring the linear combinations of two sets of variables that maximize the correlation between those two sets (Hotelling, 1992). In the present case, CCA enables the identification of the AEP **r** and the spatial pattern **a** combination that maximizes the correlation between the spatially filtered EEG data **w**^⊤^**X**, with filters **w** ∈ ℝ^*c*^ associated to the neural source **a** with **a** = **w**^⊤^cov(**X**) across *c* channels, and the common AEP **r** throughout the recording (**S***\****r**). The CCA objective then is as follows:

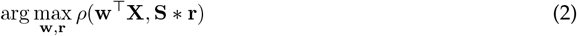

The weighting factor ***θ*** for each modality *i*, here each intensity in dB for each auditory stimulus, can be solved by finding the values of ***θ*** that maximize the correlation of the weighted AEP **r** with the brain data **X**, given a fixed **r** and **w**.

To determine an indication of the variability of the AEPs amplitude weights ***θ*** at each volume intensity, we repeat the CCA decomposition for 20 cross-validation folds—55 epochs for training the CCA, and 5 epochs to assess the model’s goodness of fit. This goodness of fit, called CCA-score, corresponds to the correlation strength between the modeled AEP from the 55 training epochs and the brain data of the 5 testing epochs. Folds were split using a shuffle split approach that samples epochs from any time point in the EEG recording.

The AEP’s amplitude weights are similar to a psychometric function: the lower the weights, the less likely the associated stimulus was perceived by the listener. Psychometric thresholds, i.e. the level at which the individual starts to perceive the stimulus, should occur at the volume intensity where the AEP’s amplitude weight starts to increase. Audiometric thresholds were automatically estimated using an algorithm that identifies the steepest increase in AEP amplitude weights, i.e., the second derivative, and places the threshold to the next volume level. This threshold estimation was exclusively employed when it was certain that the participant had perceived at least one volume intensity, as indicated by the AEP’s amplitude weights. To verify that at least one volume intensity was perceived, a one-sample t-test was calculated for each level of volume. The one-sample t-test compared the mean of the AEP’s amplitude weights across the 20 estimations from the cross-validation to a null value. The null value represents the minimum absolute AEP’s amplitude weights for the stimulus of a given frequency and presented to a given ear.

All steps and parameters in the EEG analysis pipeline (data preprocessing, modeling, and threshold estimation) were fixed before collecting the data of this study. In addition, EEG thresholds were calculated at MindAffect labs without any information about the behavioral hearing thresholds of the participants. Similarly, the behavioral thresholds were estimated at Hörzentrum Oldenburg without information about the EEG thresholds.

### 2.7 Trimmed EEG channels and recording times

An EEG test with a reduced number of electrodes and faster recording times increases the comfort and inclusiveness of patients while reducing the preparation time and testing administration costs. An 8-channel subset was identified using an extended ant colony optimization with default parameters (Izzo, 2012). Ant-colony optimization is suitable for non-linear non-convex optimization problems as it puts few assumptions on the objective function. This metaheuristic algorithm mimics the foraging behavior of an ant colony in which ants–tentative solutions–search for the shortest path–optimal solution– between the colony and a source of food. The ant-colony algorithm searched for combinations of eight channels that maximized the average CCA-score over 17 participants of other studies. Among these participants, there were hearing impaired and normal hearing participants who performed a similar EEG test (different frequencies and/or volume intensities). The participants of the present study were not included in the channel optimization process. The optimization process was run multiple times to avoid convergence to a local optimum. The optimization results were visually inspected to find which channels co-occur in good solutions and have a symmetric layout that could be easily built into a comfortable 8-channel headband. The electrodes of this solution were FC5, Fz, FC6, T7, T8, P7, P8, and POz. This 8-channel system was then validated on the current dataset to evaluate its viability on an unseen dataset. Thresholds estimated from this 8-channel EEG system were then computed and compared with 32-channel EEG and PTA thresholds using Pearson’s correlation.

In addition to a reduced EEG montage, a shorter EEG test is beneficial for patients and healthcare systems. To assess the optimal EEG test duration, audiometric thresholds were iteratively estimated starting at 2 min, ranging to 15 min in steps of 30 s, simulating real-time testing conditions. To visualize the performance over time of the EEG system, the absolute deviation between the estimated EEG thresholds and PTA thresholds was used.

### 2.8 EEG, PTA, and autoBEH threshold comparison

The correspondence between the thresholds derived from the three conditions, EEG, PTA, and autoBEH, was evaluated by pairwise comparisons between these conditions after outlier removal. At each comparison, outliers were defined as pairs of thresholds whose differences were outside of Tukey’s fences with the standard criterion of *k* = 1.5. Therefore, the number of outliers changes by comparison and affects the degrees of freedom in the Pearson correlation. The degrees of freedom are reported for each comparison.

The correspondence between the two tests was assessed by calculating fitting a linear regression between pairs of thresholds. The regression evaluated how the thresholds from two conditions correlate and if one is a good predictor of the other one with adequate slope and intercept. To that end, the metrics extracted from this regression were the Pearson’s correlation coefficient (noted as *r*_(*DF*)_, where *r* is the correlation coefficient and *DF* the degrees of freedom), the coefficient of determination *R*^2^, and the *p*-value. Additionally, the regression coefficients *β*_0_ (intercept) inform about any offset between the different types of thresholds, and *β*_1_ (slope) about the linear correspondence between the two tests.

## 3 Results

Figure 1 shows the CCA decomposition of the brain response with the spatial pattern of the AEP across EEG channels *a*, the AEP *r*, and the AEP’s amplitude weighting for each modality of the stimuli. The depicted results are an example for two participants: one hearing impaired (HI) and a control with no hearing impairment (CTRL).

**Figure 1:**
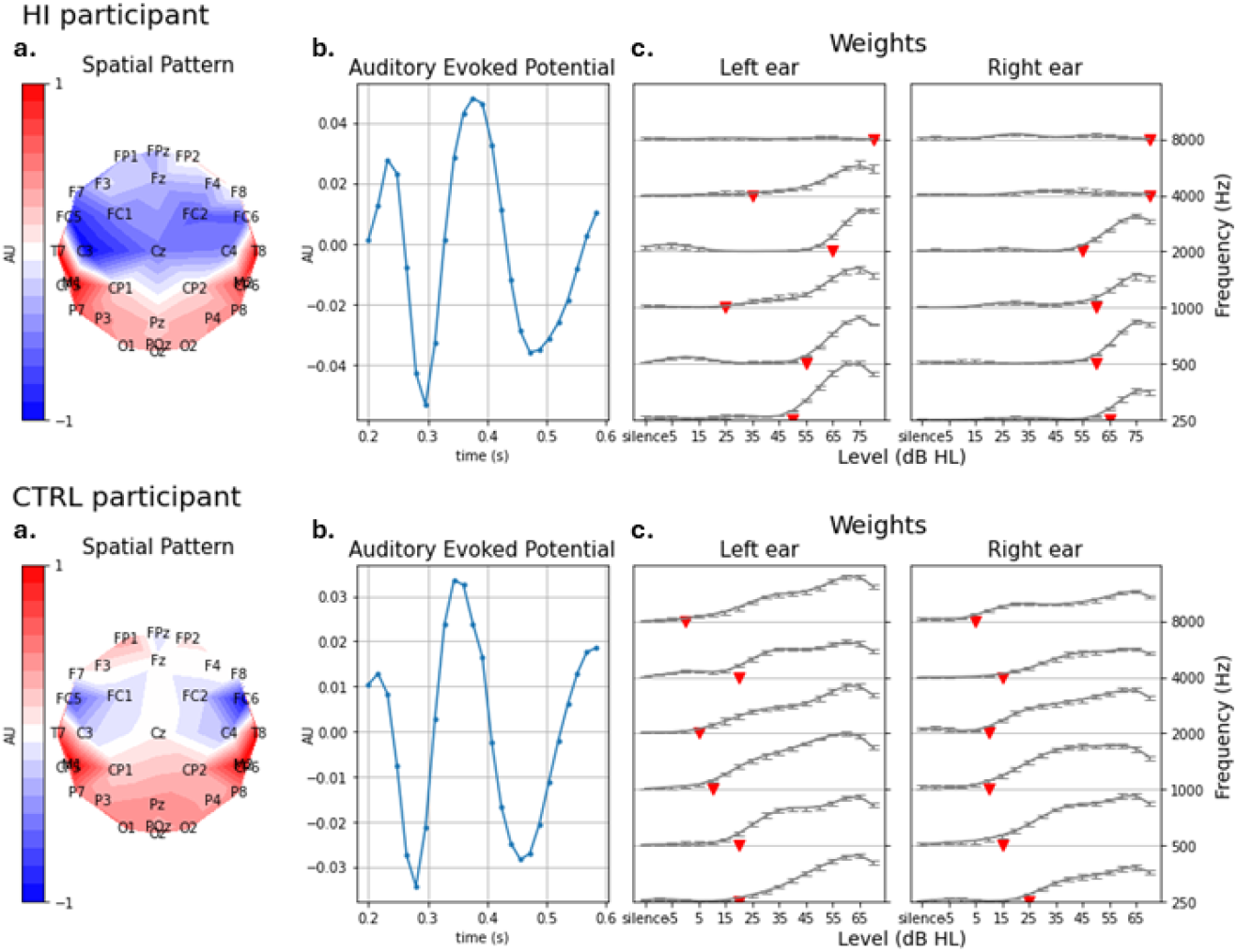
Brain responses to auditory stimuli in a hearing impaired participant (up) and normal-hearing participant (bottom): **a** panels represent the EEG spatial patterns, **a. b** panels represent the AEP, **r. c** panels show the amplitude weights, ***θ***, of the AEP for each level of stimulus volume intensity (x-axis) and for each frequency (y-axis) for the left (**c** left) and the right (**c** right) ear. Amplitude weights of the AEP, ***θ***, in the **c** panels, show the median of 20 threshold estimations with error bars representing the minimum and maximum estimates. The red inverted triangles in **c** panels show the estimated audiometric threshold for each frequency and ear

### 3.1 EEG thresholds vs PTA thresholds

Figure 2 shows strong linear correlations between EEG and PTA thresholds for 250, 500, 1000, 2000, 4000, and 8000 Hz when both groups are combined (*p*−*values <* .001). Pearson’s correlation coefficients were *r*_(28)_ = 0.80; (*β*_0_ = −14.61, *β*_1_ = 0.94), *r*_(28)_ = 0.90; (*β*_0_ = −18.46, *β*_1_ = 1.13), *r*_(25)_ = 0.90; (*β*_0_ = −10.20, *β*_1_ = 1.03), *r*_(28)_ = 0.88; (*β*_0_ = −10.98, *β*_1_ = 1.00), *r*_(25)_ = 0.89; (*β*_0_ = −6.22, *β*_1_ = 0.89), *r*_(27)_ = 0.76; (*β*_0_ = 11.69, *β*_1_ = 0.81) respectively. The linear correspondence between the two thresholds is notable through the regression slopes *β*_1_, close to 1 for all frequencies. The 4000 and 8000 Hz panels of Figure 2 present the typical increase of hearing thresholds for higher frequencies. As a result, data points are grouped into two clusters (CTRLs and HI) which might bias the linear regression fit. To mitigate this bias, a linear regression was fitted only on the HI group, see Figure 2 red regression lines.

**Figure 2:**
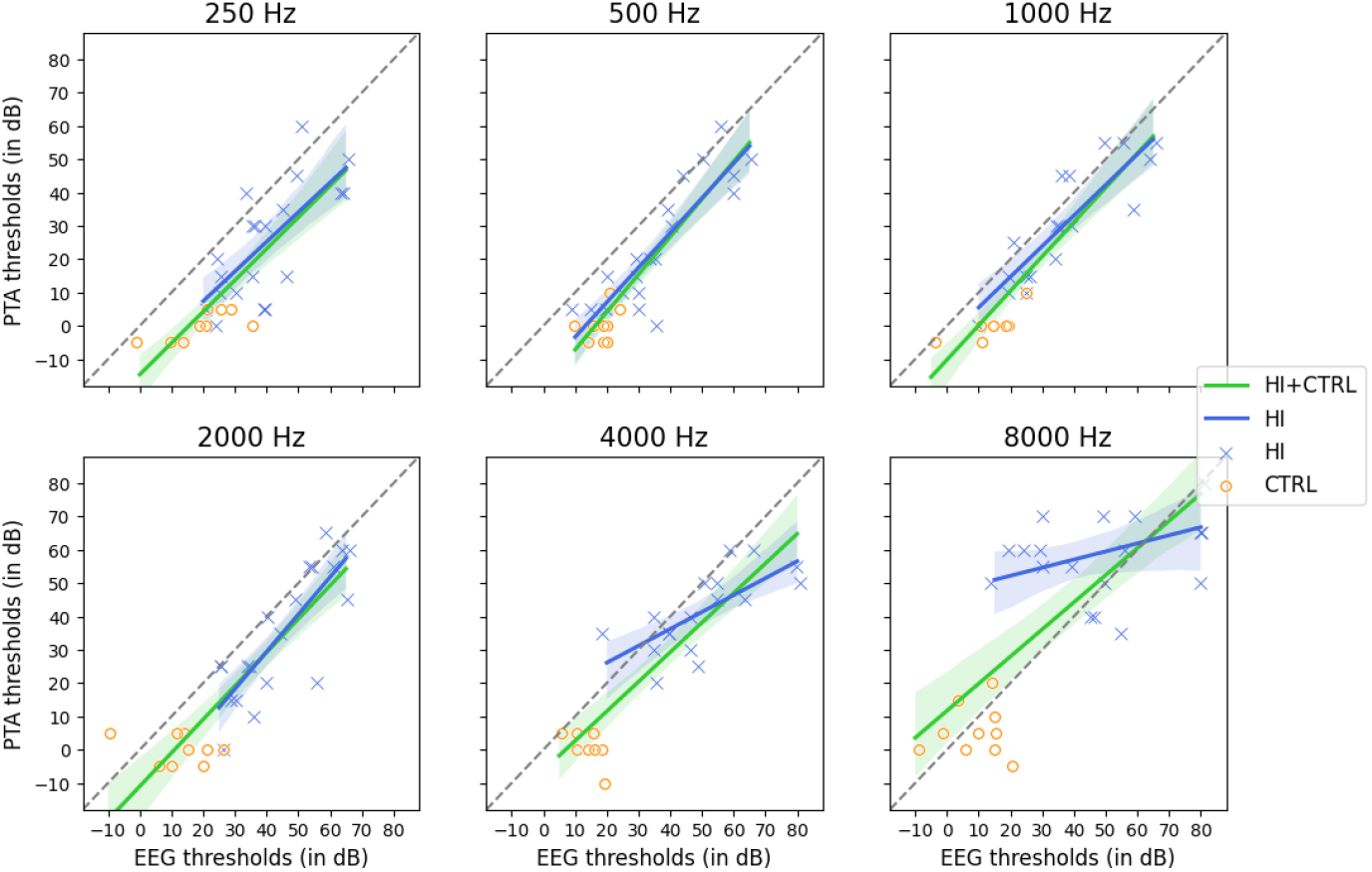
Linear regression between EEG and PTA thresholds by stimulus frequency (from top-left to bottom-right: 250 Hz, 500 Hz, 1 KHz, 2 KHz, 4 KHz, and 8 KHz) for both groups HI and CTRL (red regression line) the HI group only (blue regression line). Each marker in a panel represents a pair of thresholds derived with EEG (x-axis) and PTA (y-axis) for one subject. Given that for each frequency one threshold was estimated per ear, each participant has two associated markers. The shape and color of a marker represent the group of participants (blue cross: HI, orange circle: CTRL). The red line represents the linear regression between EEG and PTA thresholds for both HI and CTRL groups (95% confidence intervals in shaded red). The blue line represents the linear regression between EEG and PTA thresholds for the HI group only (95% confidence intervals in shaded blue). Any difference EEG - PTA that fell out of Tukey’s fences using a criterion *k* = 1.5 were removed. The gray dashed line is the diagonal, i.e., where both EEG and PTA thresholds have the same value. For the sake of the presentation, a jitter is applied to the data points on the scatter plot to prevent overlaps. This jitter is not applied to the data points used to compute the linear regressions.

Figure 2 shows a strong significant Pearson’s correlation coefficient between EEG and PTA thresholds for HI group at 250, 500, 1000, and 2000 Hz with *p* − *values <* .001 (respectively *r*_(18)_ = 0.71; (*β*_0_ = −10.38, *β*_1_ = 0.89), *r*_(18)_ = 0.87; (*β*_0_ = −13.79, *β*_1_ = 1.04), *r*_(16)_ = 0.87; (*β*_0_ = −3.71, *β*_1_ = 0.92), *r*_(18)_ = 0.84; (*β*_0_ = −15.35, *β*_1_ = 1.12)) and a moderate correlation (*p* − *values <* .05) for the 4000 Hz at *r*_(15)_ = 0.70; (*β*_0_ = 15.96, *β*_1_ = 0.51). However, no significant correlation was found at 8000 Hz (*r*_(17)_ = 0.37; (*β*_0_ = 47.33, *β*_1_ = 0.24), *p* = .24).

Figure 2 illustrates a systematic offset between PTA and EEG thresholds with different values by frequency. This offset is captured by the intercept *β*_0_. The linear regressions suggest that the offset between PTA and EEG thresholds decreases as the frequency increases. This offset could stem from the type of measurement, behavioral versus electrophysiological, and the stimulus duration. EEG testing used 30 ms stimuli while PTA used 1-3 s stimuli. To assess whether the observed offset could be attributed to the difference in stimulus duration or the nature of the thresholds, autoBEH thresholds were compared to EEG and PTA thresholds. autoBEH is a behavioral method using the same short stimuli as EEG testing. Therefore, an offset between autoBEH and PTA, but not between autoBEH and EEG would suggest that this offset is due to the stimulus duration. The results of this analysis are depicted in Figure 3

**Figure 3:**
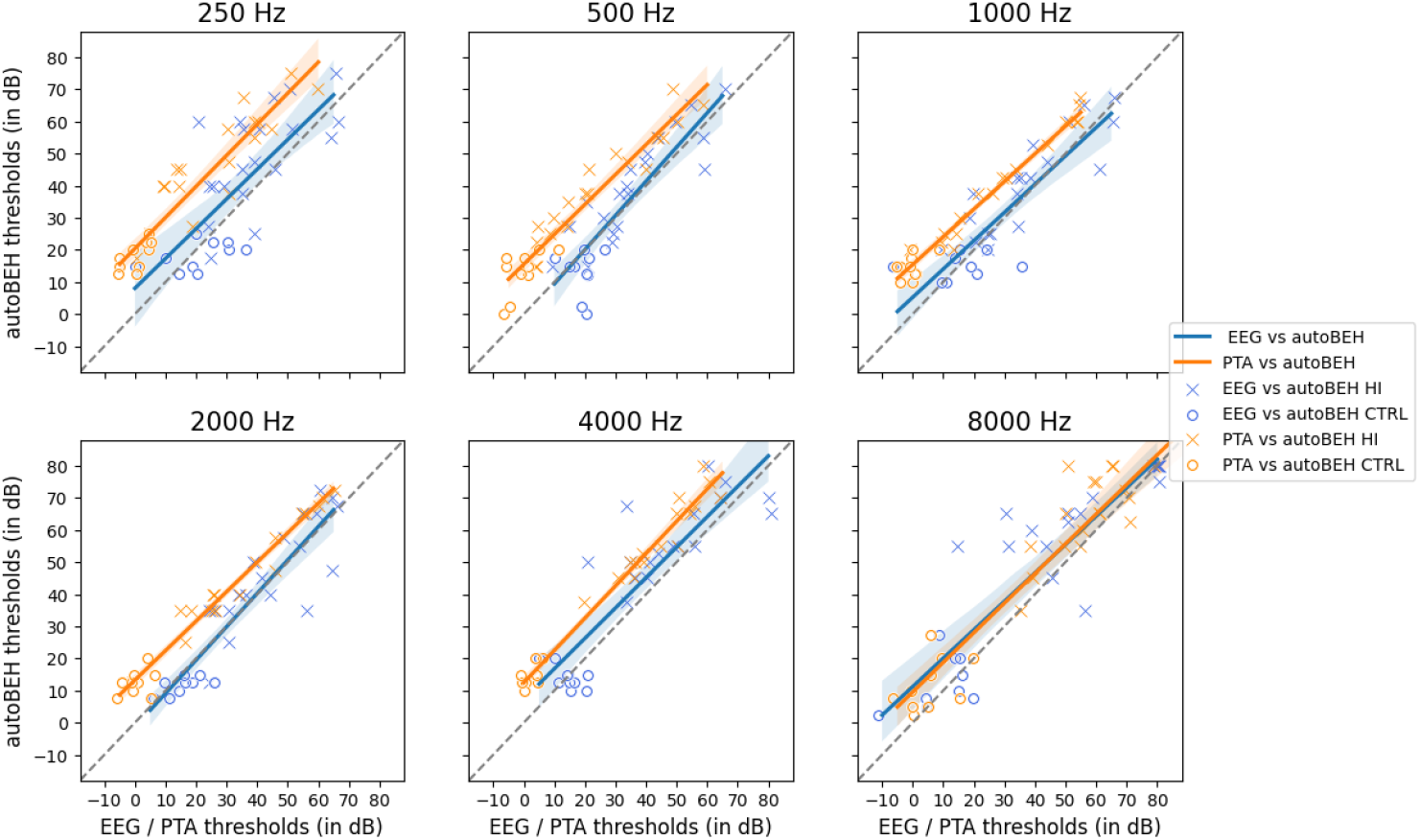
Linear regression between EEG and autoBEH (blue) and between PTA and autoBEH (orange) thresholds by stimulus frequency (from top-left to bottom-right: 250 Hz, 500 Hz, 1 KHz, 2 KHz, 4 KHz, and 8 KHz). Each marker in a panel represents a pair of thresholds derived with either PTA or EEG (x-axis) and autoBEH (y-axis) for one subject. Given that for each frequency one threshold was estimated per ear, each participant has two associated markers. The shape of a marker represents the group of participants (cross: HI, circle: CTRL). The color of a marker represents the type of comparison (blue: EEG with autoBEH, orange: PTA with autoBEH). The blue line represents the linear regression between EEG and autoBEH thresholds (95 % confidence intervals in shaded blue). The orange line represents the linear regression between PTA and autoBEH thresholds (95 % confidence interval in shaded orange). Any difference autoBEH - EEG/PTA that fell out of Tukey’s fences using a criterion *k* = 1.5 were removed.

The orange line and markers in figure 3 show the correspondence between autoBEH and PTA thresholds. Again, the results show the presence of a systematic offset by frequency which was quantified with the intercepts of the linear regressions, *β*_0_. The blue line and markers in figure 3 show the correspondence between autoBEH and EEG thresholds. Here, there is no offset between autoBEH and EEG thresholds. These findings indicate that the offset is due to the different durations of the stimulus. An analysis of variance (ANOVA) was performed to identify any factor influencing this perceptual difference characterized by the difference between behavioral thresholds estimated using short (autoBEH) and long (PTA) stimulus duration. This ANOVA considered the variable frequency, degree of hearing loss (quantified by the PTA threshold), and the interaction between these two variables as independent variables and the difference between PTA and autoBEH as the dependent variable. The results of this ANOVA show that this difference is affected by the frequency (*F* (5, 147) = 14.33, *p* − *value <* .001) and the degree of hearing loss, i.e., the PTA thresholds (*F* (1, 147) = 10.49, *p* − *value <* .01). The interaction between the two independent variables did not show any significant effect on the dependent variable (*F* (5, 147) = 0.78, *p* − *value* = .57). Those results indicate that the higher the frequency, the lower the perceptual difference. Similarly, the higher the hearing loss, the lower the perceptual difference. Therefore, these results support the hypothesis that the temporal integration, or perceptual difference, induced by the difference in the duration of the stimulus is indeed affected by frequency and hearing loss.

### 3.2 Analysis of trimmed EEG system

To evaluate the validity of the EEG system with 8 channels, we estimated audiometric thresholds exclusively from the following electrode channels: FC5, FZ, FC6, T7, T8, P7, P8, and POz. Linear associations between thresholds obtained from the 8-channel configuration and those estimated from the 32-channel configuration are depicted in Figure S1. There is a strong (*p* − *value <* .001) linear relationship between the two configurations with Pearson correlations and regression coefficients of *r*_(24)_ = 0.97; (*β*_0_ = 1.03, *β*_1_ = 1.01), *r*_(16)_ = 1.00; (*β*_0_ = 0.00, *β*_1_ = 1.00), *r*_(25)_ = 0.98; (*β*_0_ = 3.22, *β*_1_ = 0.94), *r*_(17)_ = 1.00; (*β*_0_ = 0.00, *β*_1_ = 1.00), *r*_(22)_ = 0.99; (*β*_0_ = −2.16, *β*_1_ = 1.01) and *r*_(23)_ = 0.99; (*β*_0_ = 0.56, *β*_1_ = 0.99) for 250, 500, 1000, 2000, 4000 and 8000 Hz, respectively.

To quantify the performance of the 8-channel EEG system over time and to asses shorter EEG recording times, the hearing thresholds were estimated with the 8-channel EEG configuration iteratively with increasing recording times and compared to PTA thresholds. Figure 4 shows the absolute deviation between 8-channel EEG thresholds and PTA thresholds with increasing EEG recording time. The absolute deviations between 8-channel EEG and PTA thresholds show a steep decrease until approximately 5 min of EEG recording. Then, the absolute deviation stabilizes around a persistent error for the rest of the test. A similar trend is observed for all frequencies. Next, we evaluated the correlation between the PTA and the EEG thresholds derived from 7.5 min of EEG data. The 7.5 min of EEG data were chosen because the sum of the squared error of the four frequencies at 7.5 min, as in Figure 4, corresponds with the 5^th^ percentile of the sum of the squared errors for all analysis time points.

**Figure 4:**
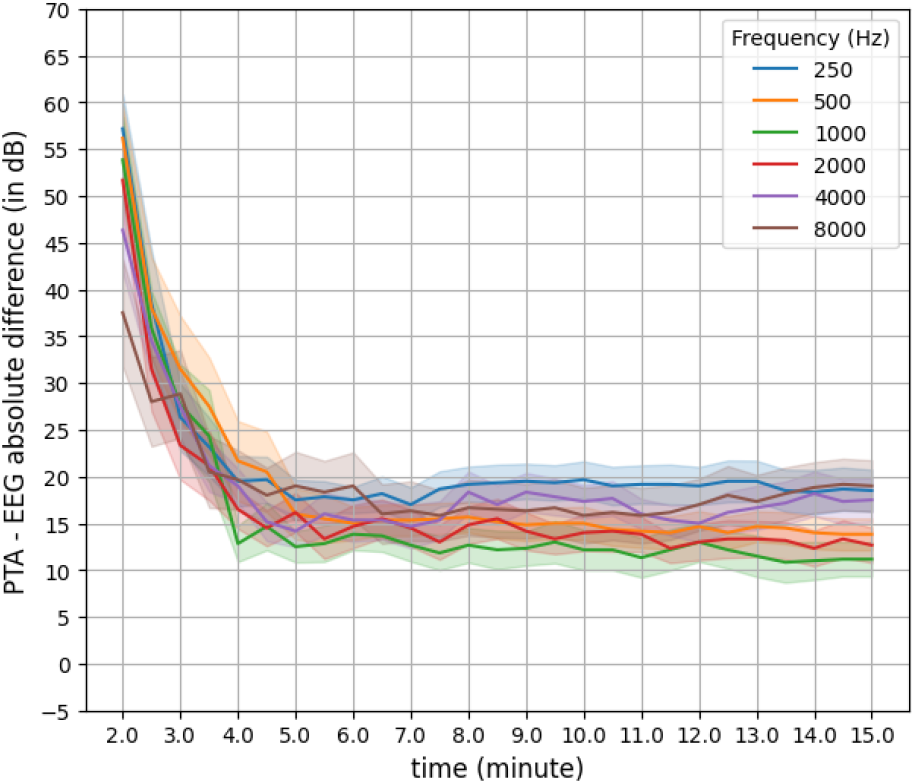
Absolute deviation between PTA and 8-channel EEG thresholds (y-axis) estimated from increasing EEG recording time in steps 30 sec (x-axis). Each color line corresponds to the absolute deviation averaged across the 15 participants and ears for one of the 6 tested frequencies. The shaded areas indicate the standard error of the mean.

**Figure 5:**
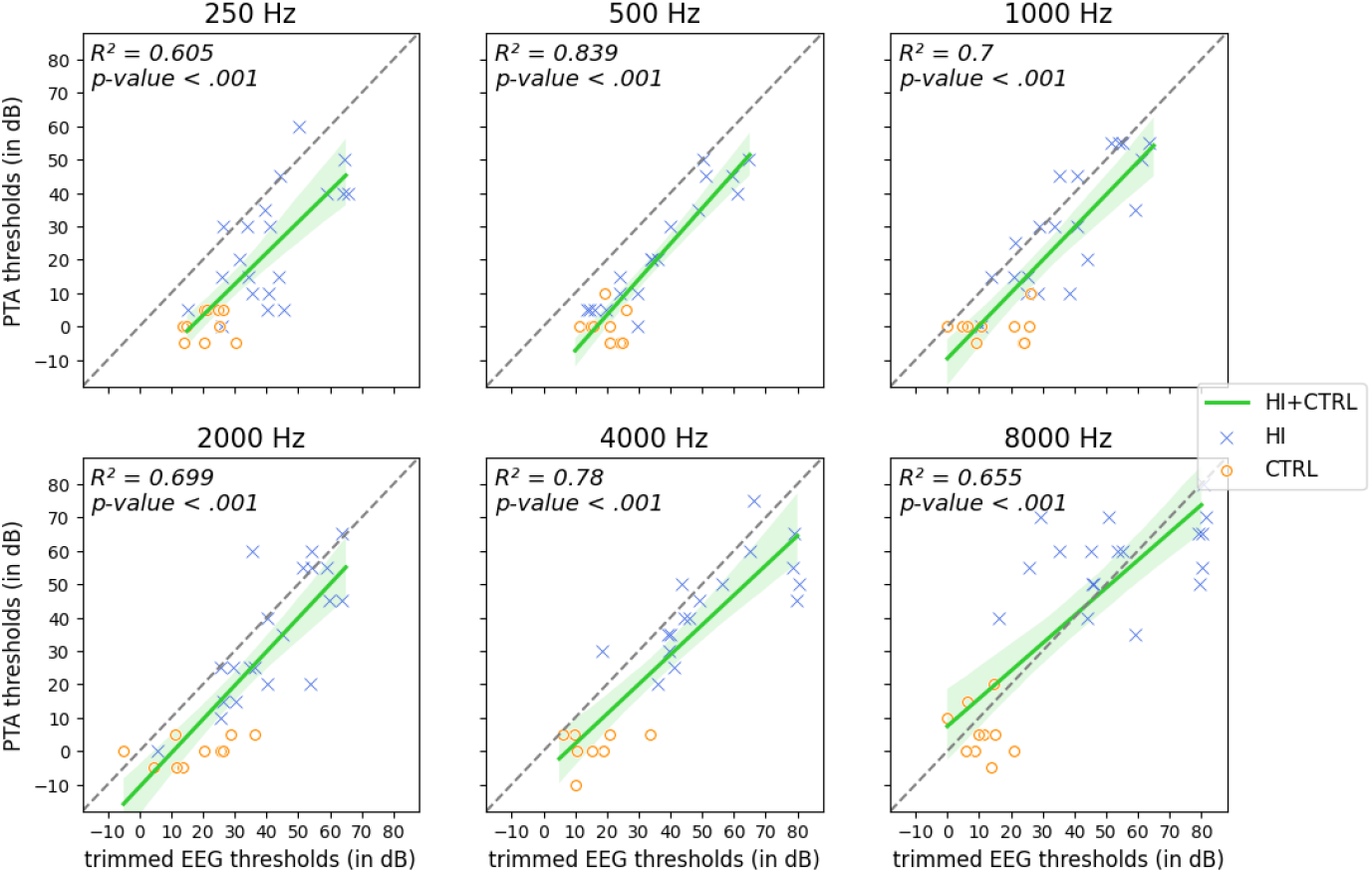
Linear regression between trimmed EEG (8-electrode and 7.5 min of EEG data) thresholds and PTA thresholds by stimulus frequency (from top-left to bottom-right: 250 Hz, 500 Hz, 1 KHz, 2 KHz, 4 KHz, and 8 KHz). Each marker in a panel represents a pair of thresholds derived from the trimmed EEG (x-axis) and PTA (y-axis) for one subject. Given that for each frequency one threshold was estimated per ear, each participant has two associated markers. The shape and color of a marker represent the group of participants (blue cross: HI, orange circle: CTRL). The green line represents the linear regression between trimmed EEG and PTA thresholds for both HI and CTRL groups (95% confidence intervals in shaded green; *R*^2^ and p-values on top-left corners of each panel). Any differences between trimmed EEG - PTA that fell out of Tukey’s fences using a criterion *k* = 1.5 were removed.

Finally, to validate the trimmed EEG system (i.e. eight electrodes and 7.5 min of EEG recording time), the Pearson correlations and regression coefficients between the trimmed EEG thresholds and the PTA thresholds were computed.

The strength and significance of the linear regression between the trimmed EEG and PTA is similar to the 32-channel EEG system after 15 min of recording (*p <* .001). Considering all frequencies, the Pearson’s correlation and regression coefficients between PTA and trimmed EEG thresholds were *r*_(171)_ = 0.82; (*β*_0_ = −9.30, *β*_1_ = 0.96). The correlations and regression coefficients between PTA and the trimmed EEG thresholds for each frequency were *r*_(28)_ = 0.78; (*β*_0_ = −15.33, *β*_1_ = 0.93), *r*_(26)_ = 0.92; (*β*_0_ = −17.91, *β*_1_ = 1.07), *r*_(28)_ = 0.84; (*β*_0_ = −9.60, *β*_1_ = 0.98), *r*_(28)_ = 0.84; (*β*_0_ = −10.77, *β*_1_ = 1.01), *r*_(24)_ = 0.88; (*β*_0_ = −6.72, *β*_1_ = 0.89), and *r*_(27)_ = 0.81; (*β*_0_ = 7.42, *β*_1_ = 0.83) for 250, 500, 1000, 2000, 4000, and 8000 Hz respectively.

## 4 Discussion

This study validates a novel EEG method to objectively estimate the audiometric thresholds for 250, 500, 1000, 2000, 4000, and 8000 Hz of each ear in 7.5 minutes with eight semi-dry EEG electrodes, as well as 15 minutes with 32-electrode semi-dry EEG electrodes. When considering all participants, EEG hearing thresholds had strong correlations and one-to-one correspondences with PTA thresholds for all frequencies. Considering the HI group only, EEG and PTA thresholds were strongly correlated for all frequencies except for 8000 Hz.

The 7.5-minute EEG test reduces drastically the common recording times of comparable objective EEG methods such as ASSR and ABR which require on average 20 and 30 minutes of recording time, respectively Sininger et al. (2018). This fast EEG audiometric test is enabled by the fast presentation rate of short pure tones and the analysis pipeline used. Furthermore, this EEG method successfully estimated all 12 hearing thresholds for all participants which is challenging for ABR and ASSR tests (Tarawneh et al., 2022; Sininger et al., 2018). The objectiveness of this new EEG method offers reliability and confidence over the estimated thresholds compared to subjective methods such as PTA. Especially when testing hard-to-test, non-collaborative, or malingering participants.

Furthermore, this study shows that the trimmed EEG system with 8 electrodes has similar performance to a 32-electrode EEG system. This compact 8-channel semi-dry EEG system allows an easier and quicker preparation, and it constitutes a more user-friendly system for technicians and patients as it does not require to apply gel in the electrodes or sticker electrodes, unlike ASSR or ABR. The analysis of the 8-channel EEG performance over time showed that after around 5 min of recording time, the estimation and quality of the model remain stable and increasing the EEG recording time does not significantly change the output of the test.

The hearing thresholds derived from EEG at 8000 Hz had a weak correspondence with PTA thresholds, unlike in all other frequencies. This exception may be explained by two factors. On the one hand, the electrophysiological measurement of the evoked responses to a pure tone of 8000 Hz has a lower amplitude than the evoked responses to other frequencies (Kuwada et al., 1986). On the other hand, elderly patients tend to have a more profound hearing loss at higher frequencies (e.g. 8000 Hz) than for lower frequencies which heavily hampers the measurement of the brain responses evoked by those high frequencies (Yang et al., 2023). The accuracy of the audiometric threshold estimation highly depends on the signal-to-noise ratio of the EEG signal of interest, i.e. the auditory evoked potential. To improve the strength of the potentials evoked by the stimuli in relation to the background EEG signals, enough auditory stimuli above the audiometric threshold should be presented. The use of a fixed range of volume intensities might fail to present enough stimuli above the hearing threshold of the participant to collect a satisfying amount of evoked potentials for modeling the evoked response which compromises the precision of threshold estimation. Taking into account the high hearing thresholds for the 8000 Hz compared to other frequencies, some of these patients would benefit from louder stimuli. However, the EEG testing should ensure that stimuli with high volume intensity do not exceed the loudness discomfort level, unless necessary. Further research and development should be performed to tackle this issue and guarantee optimal stimulation while guaranteeing patient safety and comfort.

A systematic offset between EEG and PTA thresholds was present due to a perceptual difference introduced by the distinct duration of the pure tones in the EEG and PTA tests. The literature suggests that shorter stimuli are harder to perceive than longer stimuli with identical intensity which leads to elevated audiometric thresholds in shorter stimuli. This phenomenon is regarded as the ‘temporal integration’ of a stimulus which explains the detection of a signal as a function of its intensity and duration (Gorga et al., 1984; McCreery et al., 2015; Reed et al., 2009; Watson & Gengel, 1969). Moreover, it is reported that the temporal integration decreases as the degree of hearing loss and frequency increase (McCreery et al., 2015). The current results support these statements that using different stimulus duration, i.e. 30 ms versus 1-3 s, leads to a systematic offset between the resulting thresholds. Additionally, the ANOVA showed that this systematic offset was affected by both the frequency and the degree of hearing loss, which corroborates with the literature.

To ensure the alignment between behavioral and electrophysiological methods, it is common to apply a correction factor that translates the thresholds of one system to another system. However, as of this time, there is not a single universal correction factor, and such factor differs based on the method or the system (Norrix & Velenovsky, 2017; Ghasemahmad & Farahani, 2019; Yeung & Wong, 2007; McCreery et al., 2015). To enable the translation between EEG and PTA thresholds, an additional study is required to uncover a correction factor to address this perceptual difference between long and short pure-tone. Such a study should include a large pool of participants and a large variety of variables, such as frequency, level of hearing loss, or age. This would allow the correct modeling of this perceptual difference and predict PTA threshold given an EEG threshold.

Another limitation of this study is the different headphones used for PTA and EEG (i.e., circumaural headphones and supra-aural headphones), although both headphones were calibrated according to the standards. In addition, to provide an exhaustive diagnostic tool for hearing deficits, such a system should also be able to estimate bone-conductive thresholds to differentiate between sensorineural and conductive hearing loss. Moreover, to avoid cross-hearing during bone-conduction tests, air-conductive contralateral masking should be included (Brandt & Winters, 2022). While there is no indication in this method that prevents these extra features, this study has not addressed them. A future study inspecting the feasibility of bone-conduction testing with contralateral masking will be conducted. The difference in the mean age of the two groups, HI and CTRL, could also be a confounding factor, although both groups were dissociated in some analyses without challenging the outcomes of this study. Moreover, this study evaluated the proposed EEG method on adults, but a similar EEG threshold estimation system could be applied to children. In the case of testing children with this EEG system, anesthesia might not be required because the proposed method has proved its robustness to endogenous and exogenous artifacts (Thielen et al., 2015, 2021; Martínez-Cagigal et al., 2021).

## 5 Conclusion

This study demonstrates the feasibility of estimating air-conduction hearing thresholds with a novel EEG system that uses 8 semi-dry electrodes and 7.5 minutes of EEG data. The performance of this 8-channel and 7.5-minute EEG system was not degraded in comparison with a 32-channel and 15-minute EEG system. In addition to the short recording times of this EEG system, it was conceived to be user-friendly, comfortable, and easy to set up. Therefore, the system paves the way to objectively assess the hearing thresholds of not only collaborative adults but also hard-to-test populations.

## Supplementary Information

**Table S1:**
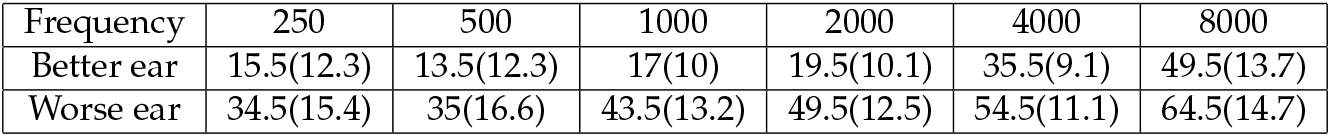
Average PTA thresholds of the HI group in dB HL (with standard deviation). Thresholds are separated between better and worse hearing ears to highlight the asymmetrical hearing loss. For 8 out of the 10 participants, the worse hearing ear is the right one and for 2 out of the 10 participants, the worse hearing ear is the left one.

| Frequency | 250 | 500 | 1000 | 2000 | 4000 | 8000 |
| --- | --- | --- | --- | --- | --- | --- |
| Better ear | 15.5(12.3) | 13.5(12.3) | 17(10) | 19.5(10.1) | 35.5(9.1) | 49.5(13.7) |
| Worse ear | 34.5(15.4) | 35(16.6) | 43.5(13.2) | 49.5(12.5) | 54.5(11.1) | 64.5(14.7) |

**Table S2:** Average PTA thresholds of the CTRL group in dB HL (with standard deviation).

| Frequency | 250 | 500 | 1000 | 2000 | 4000 | 8000 |
| --- | --- | --- | --- | --- | --- | --- |
| Both ears | 0.5(4.2) | -0.5(4.7) | -0.5(4.2) | 0(3.9) | 1(4.4) | 5.5(7.2) |

**Figure S1:**
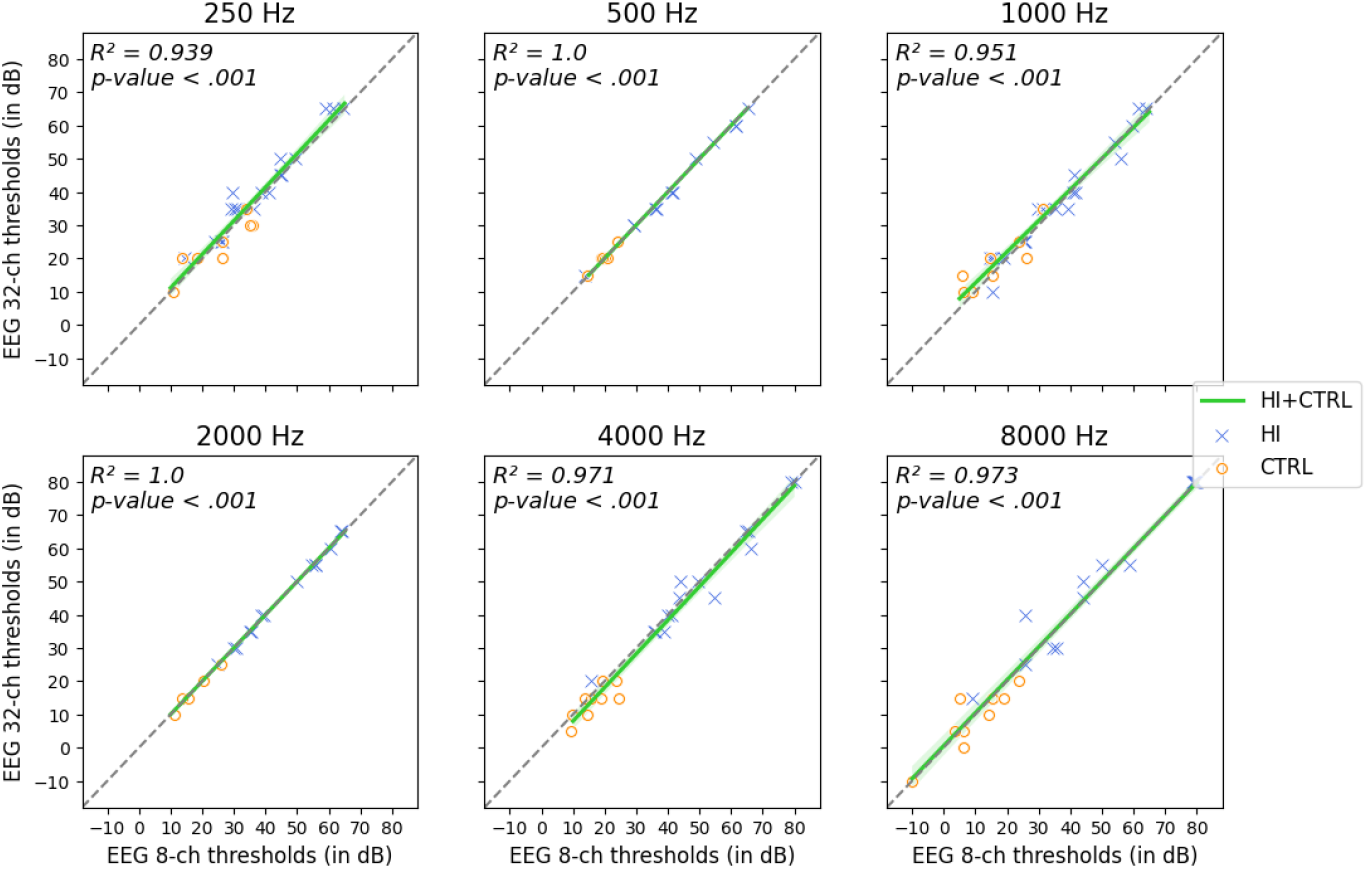
Correlation between EEG 32-channel and EEG 8-channel thresholds by stimulus frequency (from top-left to bottom-right: 250 Hz, 500 Hz, 1 KHz, 2 KHz, 4 KHz, and 8 KHz). Each dot in a panel represents a pair of thresholds derived with the 8-channel system (x-axis) and the 32-channel system (y-axis) for one subject. The shape and color of a marker represent the group of participants (blue cross: HI, orange circle: CTRL). The green line represents the linear regression between the 32-channel system and the 8-channel system thresholds (95 % confidence intervals in shaded green; *R*^2^ and p-values on top-left corners of each panel). Any difference 32-channel - 8-channel EEG testing that fell out of Tukey’s fences using a criterion *k* = 1.5 were removed. The gray dashed line is the diagonal, i.e. where both 32-channel and 8-channel EEG thresholds have the same value.

